# PLM-ICE: A Protein Language Model-Based Approach for Prediction of Ice-Nucleating and Antifreeze Proteins

**DOI:** 10.1101/2025.08.13.669380

**Authors:** Fawad Ullah, Pawel Pratyush, KC Dukka, Stephen Techtmann

## Abstract

Many microbial species have developed adaptations for coping with life in extreme cold and in particular in the cryosphere. Ice-binding proteins (IBPs) play a critical role in enabling organisms to survive in extreme cold environments. IBPs can be divided into two distinct functional classes—antifreeze proteins (AFPs) and ice-nucleation proteins (INPs). These classes have been identified based on their specific modes of interaction with ice. Here, we introduce PLM-ICE, a computational system designed to predict IBPs with high precision and sensitivity. Leveraging ESM-2 embeddings, which incorporate evolutionary and functional sequence signals more effectively than conventional embeddings, our model employs a frozen ESM-2 encoder coupled to a Multi-layer Perceptron (MLP) prediction head. The application of this architecture allows for accurate determination of AFPs and INPs, surpassing existing methods (e.g., VotePLMs-AFP) in metrics such as Matthews correlation coefficient (MCC), area under the precision-recall curve (AUPR), and area under the receiver operating characteristic curve (AUROC). Our findings indicate that PLM-ICE exhibits robust performance across broad datasets encompassing bacterial genomic sequences, highlighting its potential for wide-ranging implementation. Notably, the ability of ESM-2 to capture essential sequence patterns confers PLM-ICE with advantages in both basic research and industrial settings, where prompt and reliable identification of IBPs remains a priority. Further, the model’s strong performance underscores the broader promise of protein language model-based pipelines for decoding complex biological networks and driving innovations in cryopreservation, food technology, and climate studies. Together, these data demonstrate that PLM-ICE provides novel insight into IBP classification and stands poised to advance biotechnology applications focused on freezing tolerance and specialized temperature adaptations.

## 0.1 Introduction

Life in the cryosphere pose significant challenges to cellular structures and metabolic homeostasis, thereby challenging organisms to develop strategies that prevent or facilitate ice growth [1] and [2]. Many mechanisms for adaptation to growth at low temperatures have been identified. This broad requirement has led to two principal IBP subtypes: AFPs that inhibit ice crystal expansion and INPs that initiate freezing events at comparatively higher sub-zero temperatures. Despite seemingly opposite functions, both AFPs and INPs reflect evolutionary trajectories shaped by the demands of cold stress across diverse taxa, including fish, insects, microbes, and plants [3].

AFPs support survival in low-temperature ecosystems by selectively binding to nascent ice crystal surfaces, thereby curtailing crystal propagation. Such binding creates localized regions of freezing point depression, commonly designated as thermal hysteresis, which prevents uncontrolled ice accretion and stabilizes internal fluids [1]. Structurally, AFPs often contain repetitive amino acid motifs that align with specific planes in the ice lattice—a feature that preserves cellular membrane integrity in extreme conditions. By averting mechanical disruption and osmotic shifts attributable to freezing, AFPs safeguard organismal viability in sub-zero habitats.

In contrast, INPs coordinate water molecules along β-helical domains that are typically enriched in threonine residues, fostering organized ice growth at relatively mild freezing temperatures [2]. Through this directed nucleation process, INPs can influence atmospheric precipitation dynamics by providing templates for ice formation in clouds, thereby affecting regional weather patterns [4]. Ecologically, these proteins have considerable implications outside of their immediate physiological roles, extending to interactions in soil environments and on plant surfaces.

Historically, efforts to isolate IBPs relied on lengthy chromatographic separations, mass spectrometry, and enzymatic assays, each demanding substantial time and resources [3]. Recent advances in genomic and bioinformatic techniques have transformed IBP identification by integrating sequence features, evolutionary profiles, and structural motif data to expedite high-throughput candidate screening [5] and [6]. Tools such as iAFP [7], AFP-Pred [8], AFP-PseAAC [9], AFP-PSSM [10], TargetFreeze [11], and CryoProtect [12] have shown notable improvements in detection sensitivity over traditional lab-based methodologies [5]. This computational emphasis not only broadens the known distributions of AFPs and INPs but also refines our structural understanding of their modes of action.

Beyond their fundamental biological significance, IBPs have garnered interest for a broad array of industrial and research applications [2] and [13]. For instance, integration into cryopreservation protocols could enhance the stability of cells, tissues, or even entire organisms. Parallel opportunities exist in improving the texture and quality of frozen food products, as well as in developing anti-icing paints and coatings for aircraft and infrastructure. Ongoing advances in computational modeling of IBP–ice interactions promise to disclose novel structure–function relationships, potentially revealing new IBP variants with enhanced or specialized properties.

Yu & Lu (2011) [7] introduced a web server designed specifically for identifying antifreeze proteins, illustrating that similar sequence traits can be detected across otherwise divergent AFP structures. Their methodology employed multi-group n-peptide compositions together with genetic algorithms to reliably discriminate AFPs. Subsequently, Kandaswamy et al. (2011) [14] developed AFP-Pred, a random forest model trained on secondary structure signatures alongside physicochemical properties, yielding higher predictive accuracy relative to conventional tools including BLAST and HMM. Additional refinements emerged with AFP-PseAAC [9], which integrated pseudo amino acid composition and support vector machines, as well as AFP-PSSM [10], which harnessed evolutionary profiles—again within an SVM framework—to facilitate AFP prediction. TargetFreeze [11] further advanced feature engineering by merging the amino acid composition implicated in PseAAC and pseudo-PSSM as core inputs for SVM classification. Pratiwi et al. subsequently employed amino acid and dipeptide features in tandem with a random forest approach to establish CryoProtect [12]. The iAFP-gap-SMOTE pipeline [15] tackled uneven class distributions through a combination of feature extraction strategies and oversampling measures. Similarly, RAFP-Pred [8] stratified sequences for localized profiling, using information gain to pinpoint salient attributes for random forest-based classification. Building upon these strides, Usman et al. introduced AFP-LSE [16], leveraging an autoencoder to evaluate k-spaced amino acid pair composition. More recently, Ali et al. presented AFP-CMBPred [17], which segments position-specific scoring matrices into distinct blocks, yielding marked improvements in classification.

Protein language models (pLMs) have brought a transformative edge to protein classification, capitalizing on massive repositories of sequence data and unveiling intricate patterns that were challenging to discern with more traditional bioinformatic tools. By capturing distributed representations encompassing both the biochemical essence and evolutionary background of proteins, pLMs unlock richer insights into sequence function. VotePLMs-AFP is an exemplar of this trend, integrating embeddings from ProtT5 and ESM-1b to refine AFP identification [18]. Despite this forward momentum, AFP discovery still contends with limited data and a scarcity of definitive negative examples. In parallel, the field lacks any analogous predictive framework altogether for INPs, leaving a substantial knowledge gap. Here, to overcome these hurdles, we propose a pLM-based deep learning platform, PLM-ICE, consolidated for both AFP and INP classification. We assemble a more extensive negative set from approximately 12 million bacterial reference entries to bolster the underrepresented negative class. Distinctly, we validate our models by leveraging genome sequences from two bacterial species known to harbor both AFPs and INPs, demonstrating real-world effectiveness. Through comprehensive testing, we show that PLM-ICE is a robust and versatile system suited to the prediction of AFPs and INPs alike.

## Methods

### Dataset Curation

#### Antifreeze Proteins (AFPs) Dataset

To compile this dataset, we initially explored UniProt and identified 10,507 sequences using the keyword “Ice binding proteins.” We further examined Pfam annotations through InterPro and identified five relevant Pfam families with the following accession IDs: IPR000104 (FISH 1), IPR002353 (FISH 2), IPR006013 (FISH 3), PF11999 (Marine Bacteria), and PF05264 and PF02420 (Insect origin). Additionally, we employed protein cartography based on structural similarity to extract 1,187 antifreeze proteins (AFPs) [19], using the crystal structure and sequence of the antifreeze protein from the Antarctic sea ice bacterium *Colwellia sp. SLW05* (PDB 3WP9) as the seed structure. All identified sequences were consolidated into a single FASTA file and processed using the CD-HIT tool to eliminate redundant entries, applying a similarity cutoff of 0.5 to filter out sequences with more than 50 % similarity. This process yielded a curated positive dataset of 2,061 unique AFP sequences. For the negative class, we used a database of 12 million reference bacterial protein sequences from RefSeq complete bacterial genomes, from which 15,000 sequences were randomly selected. To ensure no overlap between the positive (AFP) and negative classes, these sequences were screened against an AFP Hidden Markov Model (HMM) and an AFP BLAST database, both developed by our team using curated AFP and ice-nucleating protein (INP) sequences. Specifically, the AFP HMM was built from 14 AFP sequences, split into 10 for training and 4 for testing, achieving 100% accuracy (4/4 test sequences predicted).

**Table 1:** Benchmark dataset composition for AFP classification.

| Dataset | AFP | RefNeg | Total |
| --- | --- | --- | --- |
| Training | 1,649 | 1,649 | 3,298 |
| Independent | 412 | 13,351 | 13,763 |

Similarly, the INP HMM was constructed from 12 INP sequences, with 8 used for training and 4 for testing, also achieving 100% accuracy. The benchmark dataset was then divided into training and independent test sets in an 80:20 ratio: 1,649 AFP sequences were randomly sampled from the 2,061 positive sequences, paired with 1,649 sequences from the 15,000 negative entries to form a balanced training set, while the remaining 412 AFPs and 13,351 negative sequences (hereafter RefNeg) comprised the independent test set for evaluating model generalizability. Table 1 and 2 provides a detailed summary of the dataset composition for AFP classification.

**Table 2:** Sequence sources and counts by Pfam accession.

| Accession | Source | Sequences | cd-hit filtered |
| --- | --- | --- | --- |
| IPR000104 | Fish 1 | 1,718 | 817 |
| IPR002353 | Fish 2 | 909 | 99 |
| IPR006013 | Fish 3 | 506 | 80 |
| PF11999 | Marine bacteria | 4,371 | 1,047 |
| PF02420 | Insect | 72 | 7 |
| PF05264 | Insect | 33 | 3 |
| PC | – | 9 | 8 |
| UniProt | – | 62 | 0 |

**Table 3:** Benchmark dataset composition for INP classification.

| Dataset | INP | RefNeg | Total |
| --- | --- | --- | --- |
| Training | 278 | 278 | 556 |
| Independent | 69 | 14,722 | 14,791 |

#### Ice Nucleation Proteins (INPs) Dataset

A similar approach was followed for Ice Nucleation Proteins (INPs) using the keyword “Ice nucleation proteins”. We retrieved 1,009 sequences from UniProt. These sequences were scanned using an Ice Nucleation Profile HMM (pHMM), which identified 444 sequences as INPs. Additionally, Pfam annotations revealed two Pfam families from bacteria, PF00818 with 288 INP sequences and PTHR31294 with 635 sequences. All INP sequences were combined into a single FASTA file, and further processed with CD-HIT using a similarity cutoff of 0.7 reduced redundancy, yielding 347 unique INP sequences. The 347 INP sequences were then divided into training and independent test sets in an 80:20 ratio, resulting in 278 sequences for training and 69 for testing. To maintain balance, 278 sequences were randomly sampled from the RefNeg dataset to serve as the negative class for training. The remaining 14,722 RefNeg sequences were retained for independent testing. A detailed summary of the INP classification dataset is shown in Table 3.

#### Sequence Embeddings

To train statistical models, FASTA sequences need to be encoded in numeric space. In this study, we utilize contextualized embeddings derived from a protein language model (LM) to encode these sequences. LMs have demonstrated great success in various protein downstream tasks, including function prediction [20], secondary structure prediction [21], and post-translational modification [22], among other [23] and [24]. Built upon the transformer architecture, these models undergo self-supervised training on extensive protein datasets, enabling them to capture the structural and functional relationships inherent in protein sequences [25]. These learned relationships are particularly valuable for characterizing AFPs and INPs as they can capture the sequence motifs, structural features, and functional patterns unique to these proteins. This ability allows LMs to identify key residues and conserved regions essential for the ice-modulating functions of AFPs and INPs, facilitating accurate predictions and deeper biological insights. In this work, we employ the state-of-the-art protein language model, ESM-2 (esm2 t6 8M UR50D variant), to encode our sequences into numerical representations [25]. ESM-2 is an encoder-only transformer model that consists of six layers, each with eight self-attention heads in its multi-head attention modules, amounting to a total of eight million parameters. For pre-training, ESM-2 was trained in a self-supervised manner on a UniRef50 dataset consisting of 49 million protein sequences.

The objective of this pretraining is masked token prediction (also referred to as Masked Language Modeling or MLM objective), where the model learns to predict masked amino acids based on their surrounding context, thereby gaining an understanding of sequence patterns and evolutionary relationships, making it highly suitable for downstream tasks such as the analysis of antifreeze and ice-nucleating proteins. Before encoding, each residue in the sequence is converted into an integer token that maps to the 20 standard amino acid alphabets, along with special tokens such as □cls□ (classification token) and padding tokens, as needed. These tokenized sequences are then processed by the ESM-2 model, which computes contextualized embeddings through its layers.

For each input sequence of length *N*, the ESM-2 model produces an embedding matrix of dimensions *L* that is the size of the final hidden layer (=360 in this case). The additional row corresponds to the embedding for the □cls□ token which is discarded, leaving an embedding matrix of dimensions *N ×* 360, representing the sequence in a token-wise manner applicable for downstream modeling.

### Proposed Architecture

The proposed architecture consists of two main components: a pLM encoder and a prediction head. The LM utilizes a pre-trained ESM-2 model with six frozen encoder layers. Features are extracted from the final hidden layer of the encoder, generating an embedding matrix with dimensions *N × L* where *L□* R^360^ and *N* is the length of the input sequence. Therefore, each residue within the sequence is represented by a feature vector of length 1 *× L*. To aggregate these features, the embedding matrix undergoes average pooling forming a protein-level representation, where the embeddings across the sequence length are aggregated using the formula:

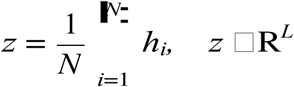

where *z* is the aggregated feature vector of length 1 *× L* (*L□* R^360^), which serves as the input to the second component, i.e., the prediction head.

The prediction head is implemented as a Multi-Layer Perceptron (MLP) consisting of two hidden layers with 64 and 32 neurons, respectively, and an output layer with a single neuron. The Rectified Linear Unit (ReLU) activation function is applied to the hidden layers, while the sigmoid activation function is used in the output layer to convert logits into probability values ranging from 0.0 to 1.0, facilitating the determination of the final prediction inferences. These inferences classify the class of the query sequence.

For AFP prediction, the model outputs whether the query sequence belongs to the AFP class or the non-AFP class. Similarly, for INP prediction, the model identifies whether the query sequence belongs to the INP class or the non-INP class. We term the overall architecture as ’PLM-ICE,’ denoting a shared design for two separate models: one tailored for AFP classification and the other for INP classification. The schematic diagram of the proposed PLM-ICE is shown in Figure **??**.

**Figure 1:**
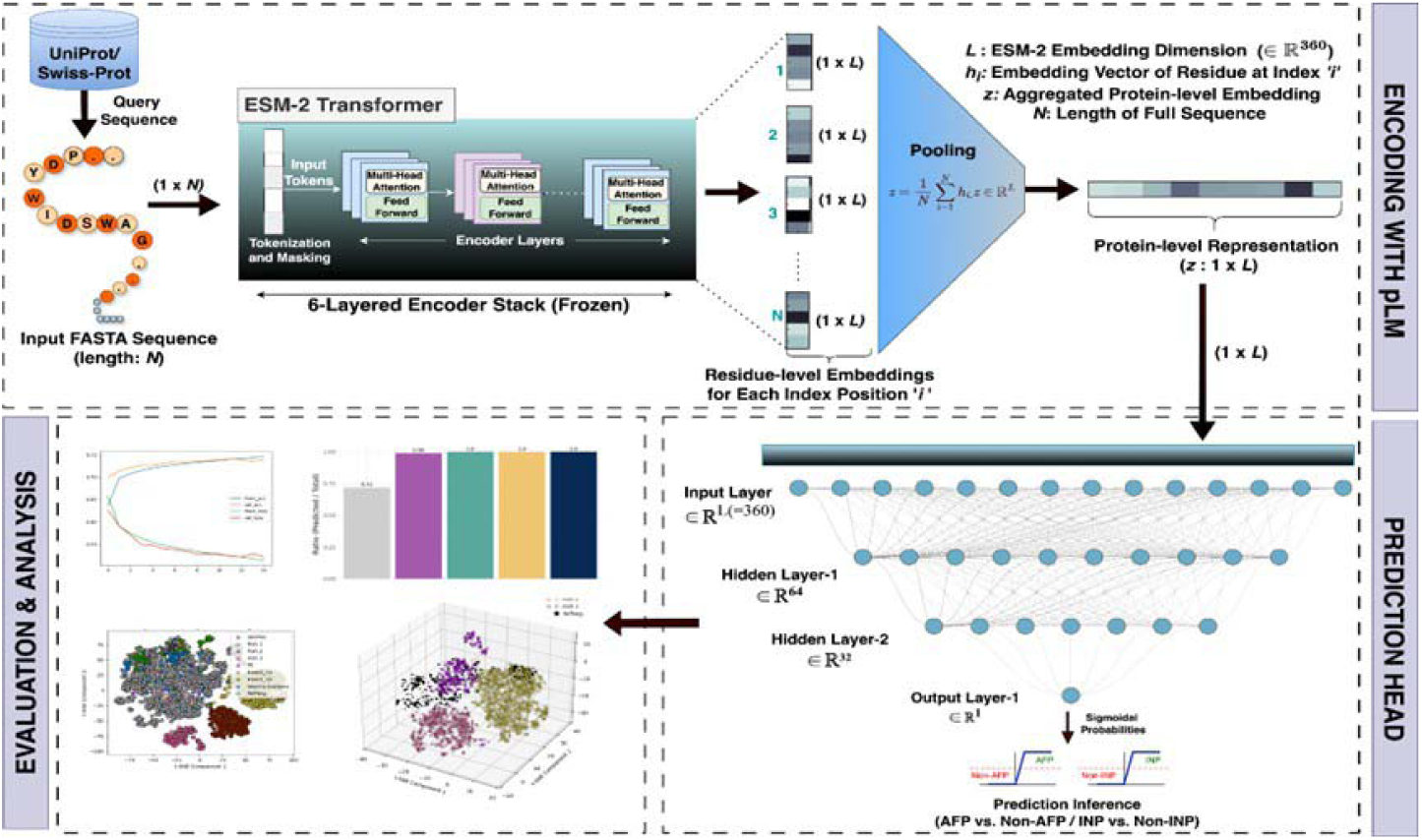
Schematic of the PLM-ICE architecture. The LM encoder processes a query sequence (length N), generating residue-level embeddings that are pooled into a protein-level representation (1×L). The MLP prediction head uses this representation for final classification (AFP/non-AFP or INP/non-INP).

### Model Training and Evaluation

In the proposed architecture, the weights of the pre-trained LM encoder layers were kept frozen during training, while the prediction heads were optimized on the datasets using the binary cross-entropy (BCE) loss or log loss function, defined mathematically as:

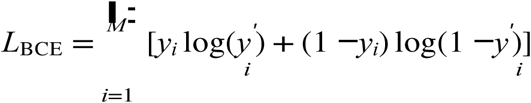

where *M* is the number of samples in the batch, *y_i_* is the true label for the *i*th sample (*y_i_□ {*0, 1*}*), and *y_i_* is the predicted probability for the positive class (*y □_i_* [0, 1]). The BCE loss was minimized using the Adam stochastic optimizer, employing an adaptive learning rate of 8 *×* 10*^−^*^5^. The model was trained for 25 epochs with a batch size of 16. Model selection and hyperparameter tuning were conducted using five-fold cross-validation on the training set, with 80% of the data used for training and 20% reserved for testing in each fold. Underfitting and overfitting were carefully monitored using accuracy/loss curves in each fold. Finally, the generalization error of the models was assessed on the independent test sets. For model evaluation, we define four fundamental metrics for binary classification:

#### True positive (TP)

Number of actual antifreeze or ice-nucleating proteins correctly predicted as positive.

#### True negative (TN)

Number of non-ice-binding proteins correctly predicted as negative.

#### False positive (FP)

Number of non-ice-binding proteins incorrectly predicted as positive (Type I error).

#### False negative (FN)

Number of ice-binding proteins incorrectly predicted as negative (Type II error).

The performance metrics were calculated as:

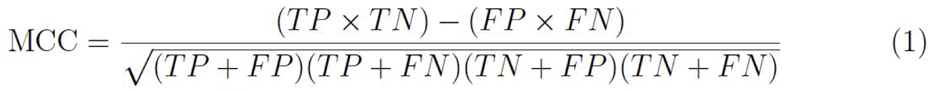

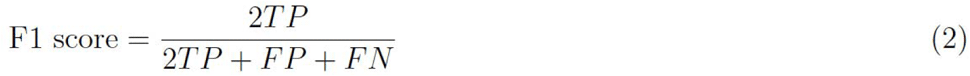

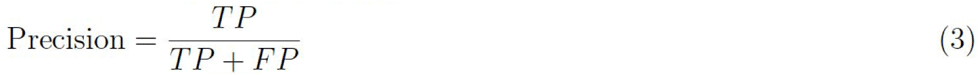

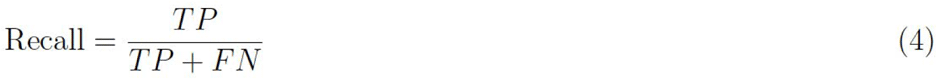

where MCC (Matthews correlation coefficient) ranges [*−*1, 1] and other metrics range [0, 1].

**Area under precision-recall curve (AUPR)** was calculated as the integral of precision with respect to recall across all decision thresholds (0.0 to 1.0).

## Results

In this section, we first present the results of the cross-validation analysis, where multiple machine learning models were evaluated to determine the optimal prediction head for both AFP (PLM-ICE-AFP) and INP (PLM-ICE-INP) tasks. This is followed by benchmarking experiments that compare the proposed PLM-ICE model against existing state-of-the-art tools, highlighting its superior performance in AFP prediction and establishing a baseline for INP prediction. Next, we present t-SNE analyses to visualize the learned feature representations and assess their discriminative power. We then evaluate the species-specific performances of the model, followed by cross-testing experiments where the PLM-ICE-AFP model is evaluated on the VotePLMs test set, and vice versa. Finally, we provide a case study evaluating our model on the reference genomes of two Pseudomonas species to validate its robustness and biological relevance.

### Cross-validation Analysis

Using five-fold cross-validation on the training sets, the optimal model was chosen for the prediction head in both AFP and INP prediction tasks. As we derived 1×L length protein-level embeddings for both tasks, we experimented with simple and shallow machine learning models. Specifically, we explored established models including Support Vector Machines (SVM), Random Forest (RF), AdaBoost (Adaptive Boosting), XGBoost (Extreme Gradient Boosting), and Multi-Layer Perceptron (MLP). Each model was subjected to a wide range of hyperparameter search spaces. Table 4 presents the comparative cross-validation results of these models for both AFP and INP tasks based on mean MCC, mean PRE, mean REC, mean F1-score, and mean AUROC. For the AFP task, several models achieved comparable performance, but the MLP model consistently outperformed others, achieving a mean MCC of 0.984, mean PRE of 0.992, mean REC of 0.992, mean F1-score of 0.992, and a competitive AUROC of 0.992, followed closely by the XGBoost model. Similarly, for the INP task, while all models demonstrated competitive performance, the MLP model stood out in most metrics, achieving a mean MCC of 0.927, mean PRE of 0.972, mean REC of 0.956, and mean F1-score of 0.962. Unlike AFP, Random Forest emerged as the second-best predictor for INP classification. Based on these results, we selected the MLP model for the prediction head in both AFP and INP tasks within the PLM-ICE architecture, as it provided the most consistent and robust performance across metrics for both tasks.

**Table 4:** Five-fold cross-validation performance of candidate models.

| Task | Model | AUPR | PRE | REC | F1 | MCC | AUROC |
| --- | --- | --- | --- | --- | --- | --- | --- |
| AFP | SVM | 0.982 | 0.983 | 0.982 | 0.982 | 0.965 | 0.998 |
|  | RF | 0.986 | 0.996 | 0.976 | 0.986 | 0.972 | 0.998 |
|  | AdaBoost | 0.979 | 0.980 | 0.978 | 0.979 | 0.958 | 0.997 |
|  | XGBoost | 0.985 | 0.979 | 0.991 | 0.985 | 0.971 | 0.998 |
|  | MLP | 0.994 | 0.992 | 0.992 | 0.992 | 0.984 | 0.992 |
| INP | SVM | 0.968 | 0.957 | 0.953 | 0.955 | 0.905 | 0.990 |
|  | RF | 0.946 | 0.960 | 0.922 | 0.941 | 0.879 | 0.990 |
|  | AdaBoost | 0.962 | 0.952 | 0.952 | 0.952 | 0.901 | 0.986 |
|  | XGBoost | 0.952 | 0.965 | 0.948 | 0.956 | 0.910 | 0.989 |
|  | MLP | 0.973 | 0.972 | 0.956 | 0.962 | 0.927 | 0.956 |

### Benchmarking AFP Prediction Performance

For AFP prediction, the proposed PLM-ICE-AFP model was benchmarked against several existing predictors, including the recently developed VotePLMs-AFP, under two scenarios to ensure a fair and robust comparison. In the first scenario, we trained and tested our model on the VotePLMs-AFP training and testing sets, comparing its performance against VotePLMs-AFP and several other predictors. These include AFP-Pred, AFP-PSSM, AFP-PseAAC, TargetFreeze, CryoProtect, RAFP-Pred, AFP-LSE, and AFP-CMBPred. The results of these predictors, as reported in the VotePLMs-AFP manuscript, were adopted for comparison. Table 5 presents the results for this scenario, where our model demonstrates superior performance, outperforming VotePLMs-AFP and all other predictors across all evaluation metrics, achieving an ACC of 0.956.

In the second scenario, we trained and tested VotePLMs-AFP on our dataset, which is larger than the VotePLMs-AFP dataset, and compared its performance against our model. Table 6 summarizes the results for this scenario, where our model consistently outperformed VotePLMs-AFP across all performance measures, achieving an ACC of 0.991. These results demonstrate the robustness and effectiveness of the proposed PLM-ICE model in AFP prediction, achieving state-of-the-art performance in both scenarios.

**Table 5:** Performance comparison of AFP prediction tools on independent test set.

| Predictor | ACC | SN | SP | Youden’s index | BACC |
| --- | --- | --- | --- | --- | --- |
| AFP-Pred | 0.833 | 0.847 | 0.823 | 0.670 | 0.835 |
| AFP-PSSM | 0.930 | 0.759 | 0.933 | 0.692 | 0.846 |
| AFP-PseAAC | 0.848 | 0.862 | 0.847 | 0.709 | 0.855 |
| TargetFreeze | 0.913 | 0.925 | 0.913 | 0.838 | 0.919 |
| CryoProtect | 0.883 | 0.873 | 0.883 | 0.756 | 0.878 |
| RAFP-pred | 0.909 | 0.840 | 0.911 | 0.751 | 0.876 |
| AFP-LSE | 0.937 | 0.867 | 0.939 | 0.806 | 0.903 |
| AFP-CMBPred | 0.917 | 0.751 | 0.920 | 0.671 | 0.836 |
| VotePLMs-AFP | 0.942 | 0.967 | 0.941 | 0.908 | 0.954 |
| PLM-ICE-AFP* | 0.956 | 0.867 | 0.958 | 0.826 | 0.913 |

**Table 6:**
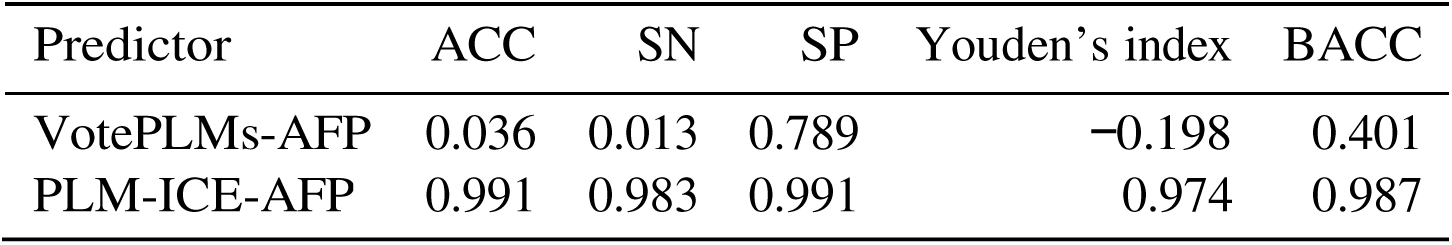
Cross-testing performance on PLM-ICE dataset.

| Predictor | ACC | SN | SP | Youden’s index | BACC |
| --- | --- | --- | --- | --- | --- |
| VotePLMs-AFP | 0.036 | 0.013 | 0.789 | −0.198 | 0.401 |
| PLM-ICE-AFP | 0.991 | 0.983 | 0.991 | 0.974 | 0.987 |

### Benchmarking INP Prediction Performance

At the time of this study, no existing computational tools were available for INP prediction. To provide a baseline comparison, we adapted the VotePLMs-AFP architecture to the INP dataset, referring to this adapted version as VotePLMs-INP for clarity. Both VotePLMs-INP and our proposed PLM-ICE-INP model were trained and tested on the INP dataset for a direct performance comparison. Table 7 presents the results of this comparison, showing that PLM-ICE-INP consistently outperforms VotePLMs-INP across all evaluation metrics. These findings underline the effective-ness of the proposed PLM-ICE model in delivering superior performance for INP prediction, further demonstrating its robustness and adaptability across both AFP and INP prediction tasks.

**Table 7:** Performance comparison of INP prediction models.

| Method | ACC | SN | SP | Youden’s index | BACC |
| --- | --- | --- | --- | --- | --- |
| VotePLMs-INP | 0.982 | 0.290 | 0.985 | 0.275 | 0.638 |
| PLM-ICE-INP | 0.987 | 0.942 | 0.987 | 0.929 | 0.964 |

### t-SNE Analysis

To assess the discriminative ability of PLM-ICE, we visualized t-SNE plots derived from the raw ESM-2 embeddings and the latent representations obtained from the hidden layers of the prediction head on the training datasets for AFP and INP independently. Figures 2a and 2b show the 2D t-SNE plots obtained from the raw embeddings for AFP and INP datasets, respectively. In these plots, some degree of separation can be observed between the positive and negative samples (AFP vs. non-AFP in Figure 2a and INP vs. non-INP in Figure 2b), reflecting the quality of the contextualized embeddings derived from ESM-2. However, there is a noticeable overlap between the positive and negative samples, suggesting the need for further refinement to achieve better discriminative power.

In Figure 2c, which visualizes the penultimate hidden layer’s representation for AFP, the separation boundary between AFP and non-AFP becomes more distinct compared to the raw embeddings. This boundary becomes even more pronounced in the final hidden layer, as shown in Figure 2d. A similar trend is observed for INP, where the separation between INP and non-INP samples improves in the penultimate hidden layer (Figure 2e) and becomes most distinct in the final hidden layer (Figure 2f). These observations demonstrate that the task-specific layers in the prediction head effectively learn features tailored to the respective tasks, enhancing the discriminative power of the PLM-ICE model.

**Figure 2:**
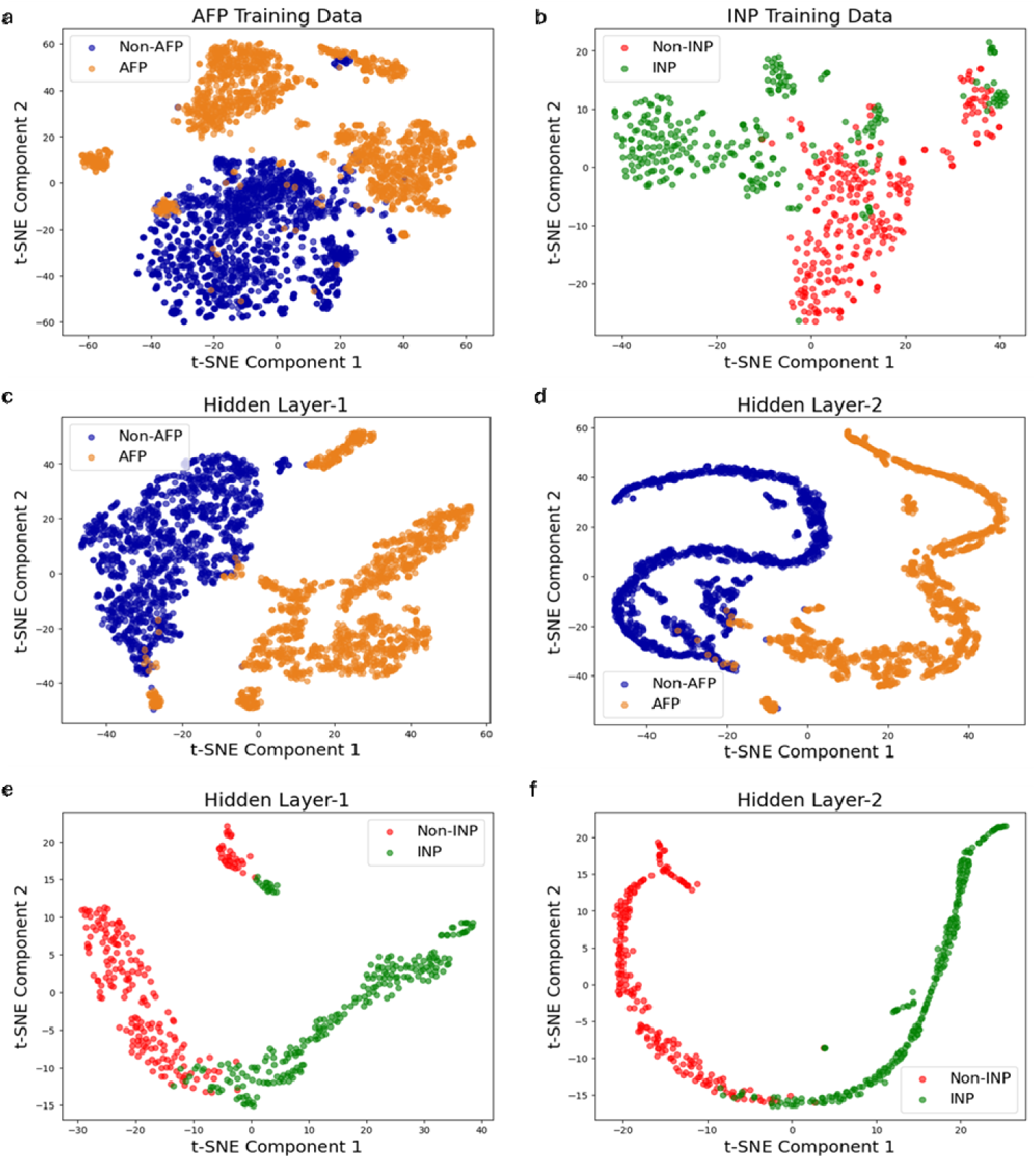
*t*-SNE visualizations of PLM-ICE model embeddings for AFP (left) and INP (right) training data. (a,d) Raw ESM-2 embeddings showing initial class separation with some overlap. (b,e) Hidden layer 1 representations. (c,f) Hidden layer 2 representations.

### Organism source-specific Analysis

The overall performance on the AFP independent test set was reported previously. To gain deeper insights into the organism source-specific performance in AFP prediction, we evaluated the model’s performance across different organism sources. Specifically, we partitioned the AFP independent test set based on organism sources (Fish 1, Fish 2, Fish 3, Insect, and Marine Bacteria) and assessed the performance of the PLM-ICE model trained on the AFP training dataset for each organism source. The evaluation metric used was the ratio, defined as the total number of correctly predicted AFP instances over the total AFP instances in the set.

The bar graph in Figure 3c illustrates the ratio proportions for each organism source AFPs that were identified by our model. Fish 1, Fish 2, and Insect achieved a perfect ratio of 100%, closely followed by Marine Bacteria with a ratio of 99%. However, the model exhibited the lowest accuracy in predicting AFPs from Fish 3. To investigate this behavior further, we visualized 2D and 3D t-SNE plots based on the raw ESM-2 embeddings . The 3D t-SNE plot in Figure 3a reveals that AFP instances from Fish 3 are positioned nearest to the negative instances, suggesting that these samples are harder to classify due to their proximity to non-AFP instances in the embedding space. Additionally, 2D t-SNE in Figure 3b further shows that the fish 3 class clustered at the bottom of the plot is a unique cluster rather than others, which are clustered in as a single group and closer to each other. This unique clustering behavior of Fish 3 underscores its distinctiveness and provides insight into the challenges associated with classifying AFPs from this organism source. The observations point to the complexity of the classification task, particularly for Fish species type 3.

### Cross-testing Analysis

We compared PLM-ICE-AFP against the existing VotePLMs-AFP architecture. This comparison involved three scenarios: 1. We replicated the original VOTE-PLM-AFP model using their architecture and training dataset (VotePLMs-AFP—VotePLMs-AFP). 2. We utilized the VotePLMs-AFP architecture but trained it using PLM-ICE-AFP dataset (VotePLMs-AFP—PLM-ICE-AFP). 3. We employed our custom architecture and training dataset (PLM-ICE-AFP—PLM-ICE-AFP).

We then assessed each model’s performance on its independent test datasets, as well as on the training and independent test datasets of the other model. The results are summarized in the (FIGURE 4). PLM-ICE-AFP—PLM-ICE-AFP generally outperforms the VOTE-PLM-AFP architecture. This suggests that our PLM-based architecture and training data are well-suited for AntiFreeze protein prediction.

**Figure 3:**
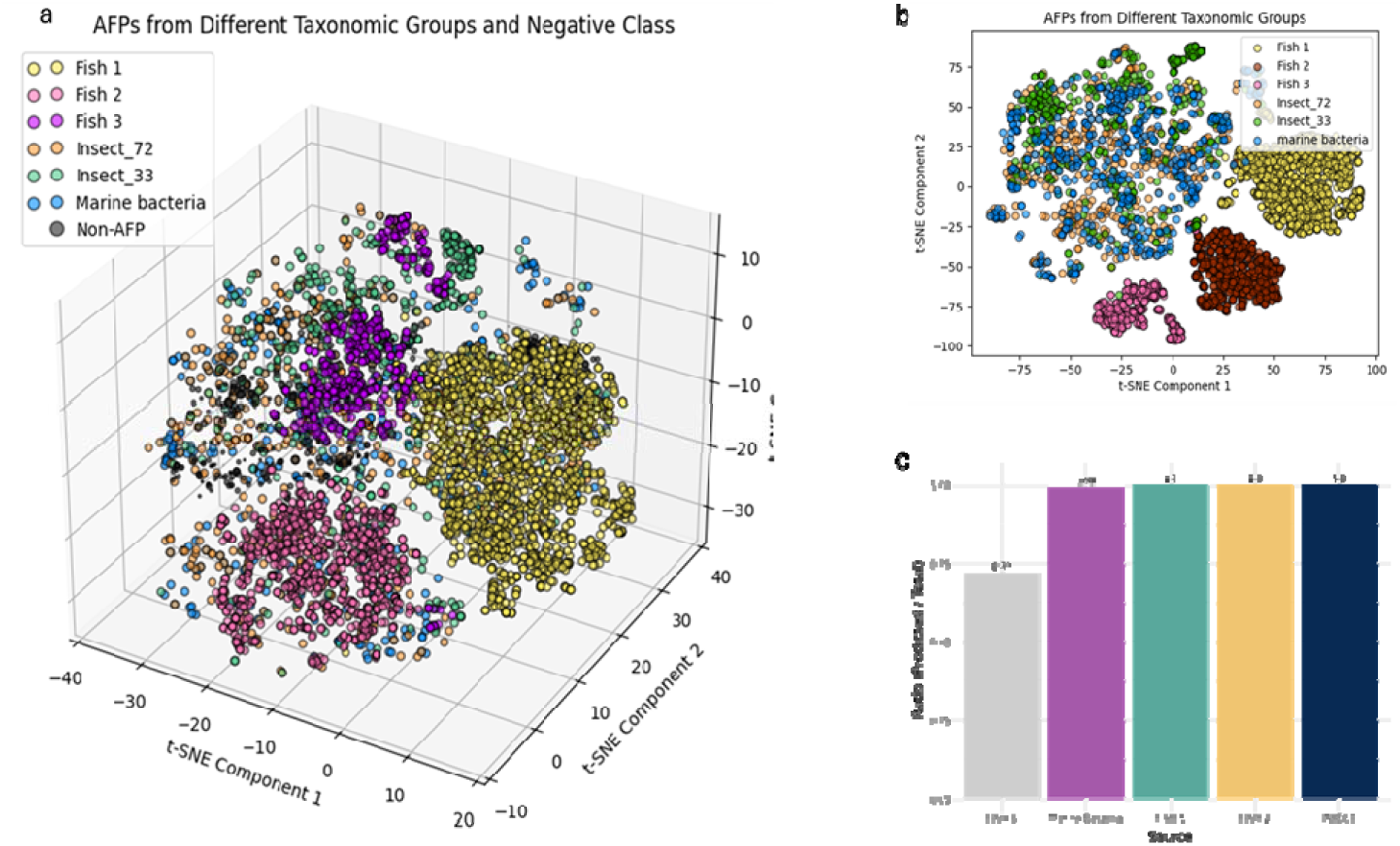
*t*-SNE visualization of AFPs across taxonomic groups versus non-AFPs. (a) 3D clusters: Fish 1, Fish 2, Fish 3, Insect_72, Insect_33, Marine bacteria, and Non-AFPs. (b) 2D separation between AFPs and non-AFPs. (c) Predicted-to-total sequence ratios per taxonomic group.

**Figure 4:**
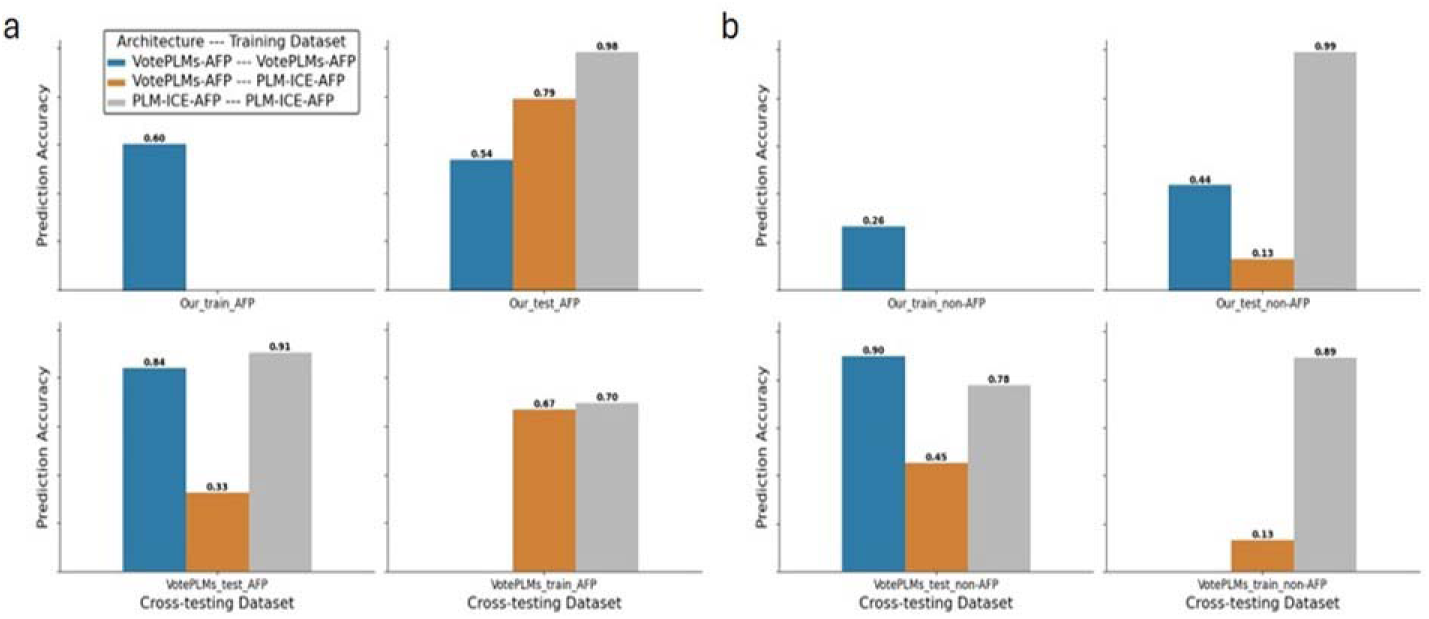
Comparison of prediction accuracy for PLM-ICE and VotePLMs-AFP models. (a) AFP sequence accuracy: training/testing on our dataset (Our train AFP, Our test AFP) and cross-testing on VotePLMs-AFP dataset (VotePLMs_train_AFP, VotePLMs_test_AFP). (b) Non-AFP sequence accuracy: training/testing on our dataset (Our train non-AFP, Our test non-AFP) and cross-testing (VotePLMs_test _non-AFP, VotePLMs_train_non-AFP). All text in figure is 8pt minimum after reduction.

**Table 8:** AFP prediction performance across *Pseudomonas* genomes.

| Genome ID | Species | Total | PLM-ICE | VotePLMs |
| --- | --- | --- | --- | --- |
| GCF_000452525.3 | *P. syringae* | 5,211 | 87 | 2,035 |
| GCF_001647715.1 | *P. antarctica* | 6,069 | 126 | 2,841 |

### Evaluation on Pseudomonas syringae and Pseudomonas antarctica

We evaluated the performance of the PLM-ICE-AFP and VotePLMs-AFP models on reference genomes obtained from NCBI for two Pseudomonas species: *Pseudomonas syringae*, which comprised a total of 5,211 proteins, and *Pseudomonas antarctica*, with 6,069 proteins. The primary motive behind this analysis was to assess the applicability of our models in real-world scenarios on real datasets. Both of these species have well-annotated genomes and contain one ice-nucleating protein (InaZ) and no annotated antifreeze proteins. The sequences from both species were screened against the two models to assess their AFP predictions. Table 8 reveals that VOTE-PLM-AFP model predicted 2,841 sequences as AFPs in the *Pseudomonas antarctica* genome. In contrast, the PLM-ICE-AFP model identified only 126 proteins as AFPs. The PLM-ICE-AFP’s prediction of 126 AFPs appears to be a more reasonable estimate, aligning with the expected biological characteristics of the species. Similarly, in the case of *Pseudomonas syringae*, the PLM-ICE-AFP model predicted 87 AFPs, while the VOTE-PLM-AFP model identified 2,035 sequences as AFPs. The PLM-ICE-AFP’s prediction is more consistent with the biological plausibility of the organism than the VOTE-PLM-AFP model. However, there are still more genes annotated as ice-nucleating proteins in these genomes than have been previously annotated. This suggests the potential for false-positive annotations even with these highly accurate models.

## Discussion

Proteins that modulate ice formation have been postulated to play fundamental roles in cold adaptation, freeze avoidance, and even climate-related processes, yet the ability to systematically identify ice-control proteins and distinguish AFPs from INPs has remained elusive. In this study, we demonstrate that leveraging advanced language modeling provides an improved framework for predicting both AFPs and INPs, thereby addressing a key gap in IBP research. Across all metrics evaluated, PLM-ICE provided a robust platform that surpassed the AFP-focused approaches and established validated benchmark for INP prediction. Previous efforts in these protein predictions have centered predominantly on AFPs which has left INPs underexplored despite the potentially broad applications for promoting ice formation in industrial, environmental, and clinical settings [26]. Here, by incorporating an expansive set of IBP sequences, we show that a single classifier can robustly differentiate AFPs, INPs, and non-IBPs. This is particularly relevant given the scarcity of experimentally validated INPs—only 347 unique sequences passed strict redundancy filters—reflecting both the limited availability of such proteins in public databases and the distinct sequence or structural patterns that may define them [27]. Our results indicate that bridging AFP and INP prediction not only provides a holistic view of IBP diversity but also offers a basis for exploring novel biological and biotechnological opportunities relevant to freeze tolerance, resource management, and cryopreservation.

One of the main challenges we faced was the class, especially for INPs, where far fewer sequences were identified relative to the voluminous negative reference set. A second layer of complexity stemmed from the fact that IBPs often show high redundancy around specific motifs, necessitating parameter tuning to reduce overfitting while preserving critical functional diversity [28]. Additionally, subtle variations in primary structure, particularly repetitive regions or organism-specific signatures explained the need for advanced methods capable of capturing nuanced sequence–structure–function relationships. These challenges mirror observations from other biological classification tasks wherein the underlying data distribution can skew algorithms away from accurately detecting minority classes [29]. When evaluated against established AFP predictors such as AFP-Pred [14], and AFP-PSSM [10], PLM-ICE consistently outperformed these tools under both cross-validation and independent test scenarios. AFP-Pred, developed by (Kandaswamy et al., 2011), uses a random forest classifier and reported an accuracy of 83.38% on an independent test dataset, with a training accuracy of 81.33%.

AFP-PSSM uses a support vector machine (SVM) with position-specific scoring matrix (PSSM) profiles and achieved an accuracy of 93.01% on a test dataset, with sensitivity of 75.89% and specificity of 93.28% [10]. PLM-ICE-AFP’s MCC reached 0.984 which highlight a near-complete segregation of AFP-positive from AFP-negative sequences. This reflects the capacity of embedding-based architectures to detect sequence determinants specific to antifreeze functionality. The previous methods such as AFP-PSSM and AFP-Pred struggle to achieve high discrimination. This could have been because of limited training dataset size, less diverse negative class selection, or their reliance on traditional machine learning architectures. PLM-ICE-AFP’s use of ESM-2 embeddings, which capture evolutionary and functional patterns, likely explains its superiority, as evidenced by its persistence in performance with added heterogeneity. Moreover, the advantage persisted when integrating newly curated AFP sequences into a larger training set which suggest that the embeddings generated by ESM-2 effectively capture relevant motifs even when additional heterogeneity is introduced.

While the INP class remains inherently data-limited, our model’s performance (MCC reaching 0.927, with high sensitivity and specificity) confirms that integral features underlying ice-nucleating capacity can be discerned from sequence data alone. Reconfiguring a prior AFP-focused tool (VotePLMs-AFP) for INP classification provided a baseline, against which PLM-ICE again showed superior metrics. This benchmark result not only demonstrates the feasibility of INP prediction but also emphasizes that ESM-2–based embeddings can identify structural repeats (e.g., β-helical folds) and amino acid motifs (often rich in threonine) that are characteristic of INPs [30].

Dimensionality reduction of the sequence embeddings via t-SNE highlighted how multi-layer perceptron–based refinement helps partition IBPs from non-IBPs in high-dimensional space. Prior to fine-tuning, the raw embeddings displayed some discernment but showed noticeable overlap between positive and negative sequences. Subsequent MLP transformations noticeably enhanced class boundaries which indicate that functional signals become more distinct when context-specific training is applied. This repositioning of borderline sequences, especially those from taxonomically diverse organisms explains the importance of additional predictive layers in refining common embedding representations. Certain fish subfamilies (Fish 3) emerged as more challenging in AFP classification, likely reflecting motif-level divergence that blurs typical antifreeze signatures. This observation resonates with parallel findings where region- or species-specific isoforms harbor atypical molecular features [31]. Future work with expanded sequence repositories may help clarify whether these variants represent novel subtypes of antifreeze proteins or functional intermediates only partially sharing conserved motifs. Practical application to the proteomes of *Pseudomonas syringae* and *Pseudomonas antarctica* buttressed the model’s reliability in large-scale searches, which is essential for functional annotation in environmental and industrial settings. Notably, PLM-ICE demonstrated a balance between high sensitivity, yet a high false-positive rates. By contrast, less stringent classifiers flagged disproportionate fractions of the proteome as AFPs, which can complicate follow-up analyses. It is possible that the high false-positive rate is due to applying PLM-ICE to large-scale genomic searches, a novel application not covered in prior studies. For instance, other models focus on sequence-based prediction without genome-wide classification, potentially missing the scale and complexity PLM-ICE addresses. This novelty could explain the high false-positive rate as the model may detect AFP-like patterns not yet annotated, and require further validation.

This work showcases that deep-learning architectures can uncover sequence determinates critical for both AFPs and INPs, paving the way for integrative IBP identification protocols. However, data limitations, particularly in the INP subset remain an obstacle to universal application. Additional experimental validation of IBP sequences, including structural and mechanistic studies of atypical or borderline variants, will be vital to improving predictive power. In this context, it is worth considering whether functional variants or partial ice-binding proteins might provide insight into evolutionary pathways connecting antifreeze and ice-nucleation activities. Our findings indicate that ESM-2–based embeddings, augmented by a dedicated MLP classifier, can successfully accommodate the stark imbalance and structural complexity characteristic of IBP prediction. The high performance in both AFP and INP detection underscores the potential utility of PLM-ICE for exploring cold-adaptive proteins on a genomic scale. As ice-binding proteins continue to gain prominence for their applications in agriculture, cryomedicine, and environmental sciences, tools such as PLM-ICE may accelerate the discovery of novel IBPs and ultimately expand our capacity to harness or mitigate ice-related processes in diverse biological and industrial systems.

## Conclusion

AFP and INPs are increasingly recognized for their essential roles in cold-adaptation processes, however our ability to accurately classify these distinct phenotypes has been limited by sparse data and shortcomings in existing tools. PLM-ICE as a unified framework leverages the contextual depth of ESM-2 embeddings to explain subtle ice-binding motifs, including β-helical folds and threonine-rich segments. This approach is particularly valuable given that many of the previously developed AFP predictors show high false-positive rates when applied to large genomic repositories, and that validated INP sequences remain scarce. Several key design strategies supported the robust performance observed. Notably, retaining the pretrained ESM-2 encoder intact minimized catastrophic forgetting, while the standalone multi-layer perceptron head provided a computationally efficient means to amplify signals indicative of IBP function. These considerations proved critical to address the class imbalance, especially for INPs, where verifiable sequence data are constrained. Further reinforcing this strategy were strict redundancy checks, along with carefully curated negative sets—a combination aimed at disambiguating borderline or partial IBP sequences that might otherwise confound large-scale annotation efforts. The near-perfect detection metrics for AFPs (MCC up to 0.984) and the high success rate in capturing INPs mark a meaningful advance over available classifiers. Beyond introducing the first computational approach for INP prediction, our findings align with the observed biodiversity of IBPs in ecologically and climatically relevant contexts, including bacterial species implicated in atmospheric ice formation. Moreover, the significantly reduced incidence of artifactual AFP calls, particularly in Pseudomonas spp. screens, points to the practicality of integrating PLM-ICE into genome-wide assessments where confidence in protein annotation is paramount. Moving forward, it will be worthwhile to explore whether these IBP-focused embeddings can be coupled to structural prediction pipelines, thus illuminating the specific residue-level interactions that underlie ice binding. In addition, expanding metagenomic analyses to underexplored or extreme habitats may reveal previously uncharacterized IBPs and help us understand how organisms thrive under subzero conditions.

## Acknowledgements

This work was supported by the Defense Advanced Research Projects Agency ReSource program cooperative agreement HR0011-24-2-0338. The views, opinions, and/or findings expressed are those of the author and should not be interpreted as representing the official views or policies of the Department of Defense or the U.S. Government.

## Notes

### Competing Interest Statement

The authors have declared no competing interest.

